# Cross-neutralization and viral fitness of SARS-CoV-2 Omicron sublineages

**DOI:** 10.1101/2022.11.08.515725

**Authors:** Hongjie Xia, Jason Yeung, Birte Kalveram, Cody J. Bills, John Yun-Chung Chen, Chaitanya Kurhade, Jing Zou, Steven G. Widen, Brian R. Mann, Rebecca Kondor, C. Todd Davis, Bin Zhou, David E. Wentworth, Xuping Xie, Pei-Yong Shi

## Abstract

The rapid evolution of SARS-CoV-2 Omicron sublineages mandates a better understanding of viral replication and cross-neutralization among these sublineages. Here we used K18-hACE2 mice and primary human airway cultures to examine the viral fitness and antigenic relationship among Omicron sublineages. In both K18-hACE2 mice and human airway cultures, Omicron sublineages exhibited a replication order of BA.5 ≥ BA.2 ≥ BA.2.12.1 > BA.1; no difference in body weight loss was observed among different sublineage-infected mice. The BA.1-, BA.2-, BA.2.12.1-, and BA.5-infected mice developed distinguisable cross-neutralizations against Omicron sublineages, but exhibited little neutralizations against the index virus (i.e., USA-WA1/2020) or the Delta variant. Surprisingly, the BA.5-infected mice developed higher neutralization activity against heterologous BA.2 and BA.2.12.1 than that against homologous BA.5; serum neutralizing titers did not always correlate with viral replication levels in infected animals. Our results revealed a distinct antigenic cartography of Omicron sublineages and support the bivalent vaccine approach.

## Introduction

Since the identification of the first Omicron sublineage B.1.1.529/BA.1 of severe acute respiratory syndrome coronavirus 2 (SARS-CoV-2) in November 2021, several other Omicron sublineages have evolved, including BA.2, BA.2.12.1, BA.2.75, BA.3, BA.4, BA.4.6, BA.5, and BF.7. Among the early Omicron sublineages (BA.1-BA.5), some circulated at low frequencies (e.g., BA.3), most likely due to their low viral fitnesses; whereas others increased in prevalence (e.g., BA.1 and BA.5), competing with and displacing some of the previous lineages over time. After the initial BA.1 sublineage surge, the BA.2 sublineage, which was more transmissible (Marc Stegger et al., 2022), became prevalent. Subsequently, derivatives of the BA.2 sublineage, such as BA.2.12.1, became predominant in the U.S. More recently, sublineages BA.4 and BA.5, which encode an identical spike (S) protein, have become prevalent and caused epidemics in different parts of the world (Cao et al., 2022; Hachmann et al., 2022; Qu et al., 2022). As of October 29, 2022 in the U.S., Omicron BA.2, BA.4, and BA.5 descendant sublineages are projected to account for 3.0%, 9.8% and 87.0%, of the total COVID-19 cases, respectively (https://covid.cdc.gov/covid-data-tracker/#variant-proportions). The selective evolution of new SARS-CoV-2 variant lineages is mainly driven by two forces: (i) evasion of immunity elicited from infection and/or vaccination and (ii) improved viral replication or transmission in the human host (Frederik Plesner Lyngse et al., 2022; Liu et al., 2022b; Sandile Cele, 2021). Compared with all the previous variants of concern (*i.e*., Alpha, Beta, Gamma, and Delta), Omicron variants exhibited the greatest immune evasion (Evans et al., 2022; Kurhade et al., 2022a; Liu et al., 2021b; Liu et al., 2021d; Liu et al., 2021e; Sandile Cele, 2021; Wang et al., 2022). Among the Omicron sublineages, BA.5 is least susceptible to antibody neutralization when tested against previous variant-infected convalescent sera or vaccinated sera (Kurhade et al., 2022b; Xie et al., 2022; Zou et al., 2022b). Compared with the index SARS-CoV-2 isolated in early 2020, the S proteins of different Omicron sublineages have accumulated more than 30 amino acid substitutions (Viana et al., 2022). Thus, for future vaccine development, it is important to understand (i) the antigenic relationship and (ii) the relative viral replication among different Omicron sublineages. In this study, we used a mouse model and a primary human airway cultures to address these two important questions.

## Results

### Experimental rationale

For studying the antigenic relationship among different Omicron sublineages, it was challenging to obtain human sera from individuals who were infected with distinct Omicron sublineages without prior SARS-CoV-2 infection or vaccination. To overcome this challenge, we used a panel of sera collected from mice, which were infected with distinct Omicron sublineages, to examine their antigenic relationship. We infected K18-hACE2 mice with recombinant Omicron BA.1, BA.2, BA.2.12.1, or BA.5 (**Fig. 1A**). The selection of these four sublineages was based on their roles in causing Omicron waves after the initial BA.1 emergence. The mouse convalescent sera were examined for their neutralization titers against five Omicron sublineages (BA.1, BA.2, BA.2.12.1, BA.4.6, and BA.5), a Delta variant, and USA-WA1/2020, the reference index SARS-CoV-2. The resulting neutralization titers enable development of antigenic cartography to illustrate antigenic relationships among various Omicron sublineages. Additionally, we could also compare the viral replication of various Omicron sublineages in K18-hACE2 receptor transgenic mice and in primary human airway epithelial (HAE) cultures. Furthermore, the K18-hACE2 mouse model data allow comparison of pathogenicities among Omicron sublineages as well as to analyze the relationship between viral replication levels and neutralization activities.

**Figure 1.**
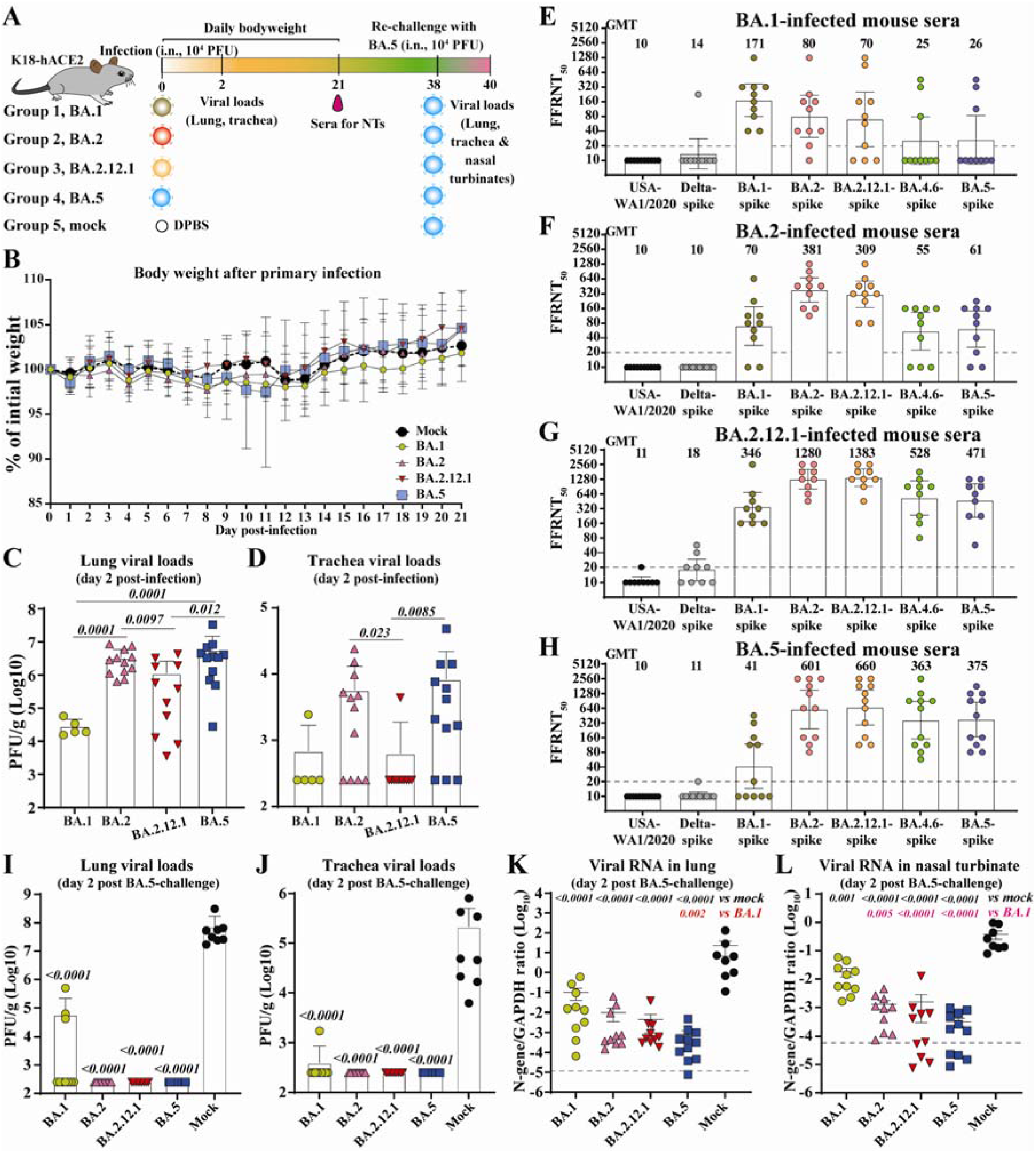
Cross-neutralization of SARS-CoV-2 variants elicited by BA.1, BA.2, BA.2.12.1, or BA.5 infection. **(A)** Experimental scheme. 8-10-week-old K18-hACE2 mice were infected via intra nasal (i.n.) route with infectious clone derived BA.1, BA.2, BA.2.12.1 or BA.5. On day 21 post-infection, mice sera were collected and cross-neutralization against SARS-CoV-2 variants was measured by FFRNT. On day 38 post-primary infection, all mice were rechallenged with BA.5. Two days after the challenge, mice were euthanized, and tissue viral loads were quantified. (B) Mouse body weight after primary infection. Daily body weight was normalized to the initial body weight. Data are presented as mean ± standard deviation (SD). (C) Lung viral loads after day 2 post-infection. (D) Trachea viral loads after day 2 post-infection. (E) FFRNT_50_s of BA.1-infected mouse sera. The dashed line indicates the limit of detection (1:20 dilution). The bar heights and the numbers above indicate geometric means of neutralizing titers (GMT). The whiskers indicate 95% CI. The Wilcoxon matched-pairs signed-rank test was performed for group comparison of GMTs. The *p* values between GMTs against BA.1-spike and USA-WA1/2020, Delta-spike, BA.2-spike, BA.2.12.1-spike, BA.4.6-spike and BA.5-spike are 0.002, 0.002, 0.082, 0.30, 0.049, 0.065, respectively. (F) FFRNT_50_s of BA.2-infected mouse sera. The *p* values between GMTs against BA.2-spike and USA-WA1/2020, Delta-spike, BA.1-spike, BA.2.12.1-spike, BA.4.6-spike and BA.5-spike are 0.002, 0.002, 0.002, 0.10, 0.002, 0.002, respectively. (G) FFRNT_50_s of BA.2.12.1-infected mouse sera. The *p* values between GMTs against BA.2.12.1-spike and USA-WA1/2020, Delta-spike, BA.1-spike, BA.2-spike, BA.4.6-spike and BA.5-spike are 0.004, 0.004, 0.008, 0.63, 0.016, 0.008, respectively. (H) FFRNT_50_s of BA.5-infected mouse sera. The *p* values between GMTs against BA.5-spike and USA-WA1/2020, Delta-spike, BA.1-spike, BA.2-spike, BA.2.12.1-spike, and BA.4.6-spike are 0.001, 0.001, 0.001, 0.004, 0.008, 0.84, respectively. (I) Lung viral loads at day 2 post-challenge. (J) Trachea viral loads at day 2 post-challenge. (K) Viral RNA in the lung on day 2 post-challenge. (L) Viral RNA in nasal turbinates at day 2 post-challenge. The dashed line in (K-L) shows the cutoff as determined from the uninfected mouse samples. Data in (C, D, I-L) show mean ± standard deviation (SD). *p* values shown as italic numbers in (C, D, I-L) were calculated using one-way ANOVA with Tukey’s multiple comparisons test. *p*<0.05, statistically significant.

### Viral replication in K18-hACE2 mice

We constructed infectious cDNA clones for four viruses representative of Omicron sublineages: BA.1 (GISAID EPI_ISL_6640916), BA.2 (GISAID EPI_ISL_11253924.1), BA.2.12.1 (GISAID EPI_ISL_12115772), or BA.5 (GISAID EPI_ISL_11542604). The Omicron recombinant viruses were prepared using a well-established protocol (Xie et al., 2021; Xie et al., 2020). All four recombinant viruses were sequenced to ensure no undesired mutations. To compare viral replication and pathogenesis, we intranasally inoculated 8- to 10-week-old K18-hACE2 mice with 10^4^ PFU of each of the four recombinant viruses (**Fig. 1A**). No significant weight loss was observed for most of the groups of mice (**Fig. 1B**), except one mouse from the BA.5 group lost >20% body weight on day 11 post-infection and had to be humanely euthanized following an approved IACUC protocol. Different viral loads were detected in the respiratory tracts for different sublineage groups. The infectious viral loads in the lungs were in the order of BA.5 ≈ BA.2 > BA.2.12.1 > BA.1 (**Fig. 1C**), while the viral loads in the trachea were in the order of BA.5 ≥ BA.2 > BA.2.12.1 ≈ BA.1 (**Fig. 1D**). Thus, the rank order of viral replication efficiency in the K18-hACE2 model was in the order of BA.5 ≥ BA.2 > BA.2.12.1 ≥ BA.1.

### Viral replication in primary human airway epithelial (HAE) cultures

To substantiate the K18-hACE2 mouse results, we compared the replication kinetics of BA.1, BA.2, BA.2.12.1, and BA.5 on HAE culture. The HAE culture has been reliably used to study viral replication of SARS-CoV-2 variants, yielding results that recapitulate viral fitness in humans (Liu et al., 2022a). Competition experiments were used to directly compare replication kinetics of two sublineage viruses in HAE cultures (**Fig. 2A**). A mixture of two viruses was used to infect HAE cultures. The relative replication of the two viruses was quantified by next-generation sequencing of the viral RNA population collected on different days post-infection. This competition assay has been reliably used to compare viral replication between variant viruses (Liu et al., 2021a; Liu et al., 2021c; Plante et al., 2021; Shan et al., 2020). Our competition results showed that (i) BA.2 replicated more efficiently than BA.1 (**Fig. 2B-C**); (ii) BA.2 replicated slightly more efficiently than BA.2.12.1 (**Fig. 2D-E**); and (iii) BA.5 replicated more efficiently than BA.2 (**Fig. 2F-G**) and BA.2.12.1 (**Fig. 2H-I**). Overall, the HAE results suggest the viral replication efficiency in the order of BA.5 > BA.2 ≥ BA.2.12.1 > BA.1.

**Figure 2.**
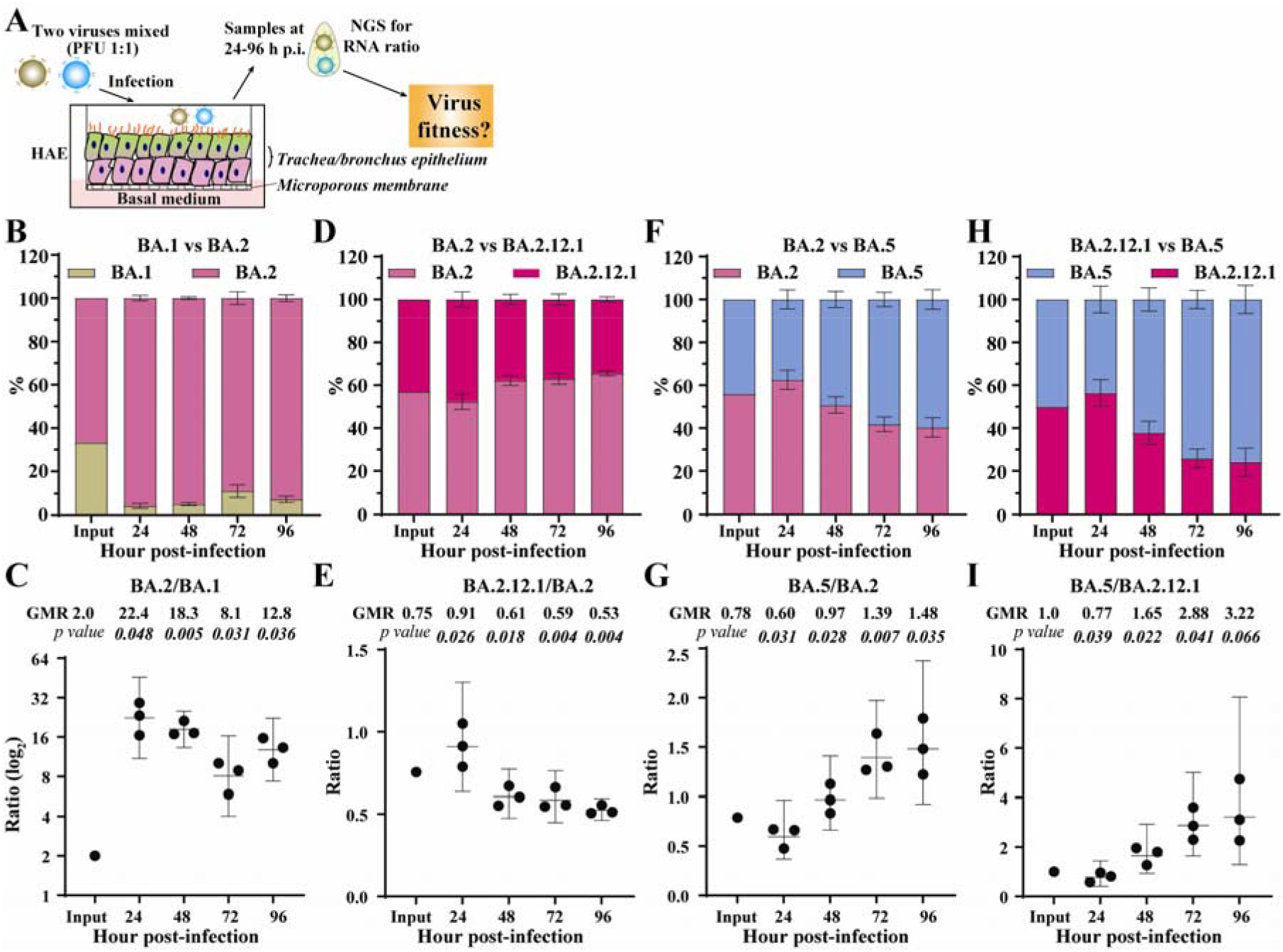
Competition of Omicron sublineages in HAE. (A) Diagram of competition experiment. Human airway epithelium was infected with a mixture of two viruses (equal PFU). At 24-96 h post-infection, extracellular viral RNA populations were determined by next-generation sequence (NGS). The viral RNA population in the inoculum measured by NGS was indicated as input. (B) The percentage of viral RNA in BA.1 and BA.2 infected HAE. (C) Scatter plot of the ratio of BA.2 to BA.1 RNA from infected HAE. (D) The percentage of viral RNA in BA.2 and BA.2.12.1 infected HAE. (E) Scatter plot of the ratio of BA.2.12.1 to BA.2 RNA from infected HAE. (F) The percentage of viral RNA in BA.2 and BA.5 infected HAE. (G) Scatter plot of the ratio of BA.5 to BA.2 RNA from infected HAE. (H) The percentage of viral RNA in BA.2.12.1 and BA.5 infected HAE. (I) Scatter plot of the ratio of BA.5 to BA.2.12.1 RNA from infected HAE. The mean ± standard deviations (SD) are shown in B, D, F, and H. The geometric ratio (GMR) and 95% confidence intervals (indicated as error bars) are shown in C, E, G, I. P values are calculated as the coefficient of each linear regression analysis of RNA ratio at a given time-point versus input RNA ratio.

### Cross-neutralization between different Omicron sublineages

To examine cross-neutralization and antigenic relationships among different Omicron sublineages, we collected sera from BA.1-, BA.2-, BA.2.12.1-, or BA.5-infected K18-hACE2 mice on day 21 post-infection (**Fig. 1A**). Each serum was measured for neutralizing antibody titers against homologous and heterologous variant-spike viruses, including the four Omicron sublineages (BA.1, BA.2, BA.2.12.1, and BA.5), a recent Omicron sublineage (BA.4.6), the index USA-WA1/2020, and a Delta variant (**Figs. 1E-H**). To specifically study the impact of amino acid changes in the spike and increase the throughput of the neutralization assay, we engineered the full-length spike gene from each variant into the backbone of an attenuated USA-WA1/2020 that lacked open-read-frame-7 (ORF7), an accessory gene that is not essential for viral replication *in vitro* but facilitates replication *in vivo* (Johnson et al., 2022; Liu et al., 2022d). The ORF7 gene was replaced with the mNeonGreen (mNG) reporter gene (Muruato et al., 2020; Xie et al., 2020). The resulting variant-spike USA-WA1/2020 mNG viruses (**Fig. S1**) were used for a fluorescent focus reduction neutralization test (FFRNT). The FFRNT assay has been proven reliable for high-throughput neutralization measurement (Liu et al., 2022c; Zou et al., 2022a).

The FFRNT results revealed distinct cross-neutralization profiles for BA.1- (**Fig 1E**), BA.2.- (**Fig. 1F**), BA.2.12.1- (**Fig. 1G**), and BA.5-infected mouse convalescent sera (**Fig. 1H**). Three conclusions could be drawn from the cross-neutralization data. First, the cross-neutralization titers against various Omicron sublineages were significantly higher than those against USA-WA1/2020 or Delta variant. The sera from BA.1-, BA.2.-, BA.2.12.1-, or BA.5-infected mice exhibited no or almost no neutralizing activities against the original USA-WA1/2020 or Delta variant, with FFRNT_50_s <20 (**Figs. 1E-H**), indicating the antigenic divergence among these variants. Second, sera from different Omicron sublineage infections showed distinct neutralizing patterns against homologous and heterologous variant-spike SARS-CoV-2. BA.1-, BA.2.-, and BA.2.12.1-infected animals developed higher neutralizing titers against homologous variants than those against hetereologous variants (**Figs. 1E-G**), illustrating that they are antigenicially distinguishable from each other. In contrast, BA.5-infected mice developed higher neutralization titers against heterologous BA.2 and BA.2.12.1 variants than against the homologous BA.5 variant (**Fig. 1H**). Third, serum-neutralizing titers did not necessarily correlate with viral replication levels in the respiratory tracts of infected animals. This is exemplified by BA.2.12.1 which replicated less efficiently than BA.2 and BA.5 in both lungs and tracheas (**Figs. 1C & D**), but elicited higher neutralizing titers than the BA.2 and BA.5 viruses(Compare **Fig. 1G** with **Figs. 1F & H**). This result also suggests that BA.2.12.1 may be more immunogenic than BA.1, BA.2., and BA.5. The higher immunogenicity of BA.2.12.1 could be determined by mutations inside and/or outside the S gene.

### Cross-protection of BA.1-, BA.2-, BA.2.12.1-, or BA.5-infected mice against Omicron BA.5 challenge

BA.5 SARS-CoV-2 was used to challenge K18-hACE2 mice that were immunized by infection with BA.1, BA.2, BA.2.12.1, or BA.5 variants for two reasons: (i) BA.5 was the most prevalent sublineage around the world at the time of the experiment and (ii) BA.5 was the least neutralized sublineage among the tested Omicron sublineages by the BA.1- and BA.2-infected mouse sera. On day 38 after the initial BA.1, BA.2, BA.2.12.1, or BA.5 infection/immunization, the mice were intranasally inoculated with 10^4^ PFU of recombinant BA.5 SARS-CoV-2 (**Fig. 1A**). The mock-immunized group developed mean titers of 7.3×10^7^ PFU/g and 2.1×10^5^ PFU/g of infectious BA.5 SARS-CoV-2 on day 2 post-challenge in lungs (**Fig. 1I**) and tracheas (**Fig. 1J**), respectively. In contrast, only a few BA.1-immunized mice developed low titers of infectious BA.5 virus in the lungs (**Fig. 1I**) and tracheas (**Fig. 1J**). No infectious viruses were detected in the respiratory tracts of the BA.2-, BA.2.12.1-, or BA.5-immunized animals after BA.5 challenge (**Fig. 1I-J**). To increase the detection sensitivity, we also performed quantitative RT-PCR to quantify the viral RNA in the respiratory tracts after the challenge. Compared with the mock-immunized group, the BA.1-, BA.2-, BA.2.12.1-, and BA.5-infected groups showed 223-, 2,253-, 4,816-, and 27,702-fold reduction of viral RNA in lungs (**Fig. 1K**) and 21-, 290-, 242-, and 1,157-fold reduction in nasal turbinates (**Fig. 1L**), respectively, after the BA.5 challenge. The decreased levels of BA.5 RNA reversely correlated with the neutralizing titers against BA.5. Specifically, the BA.1- and BA.2-infected sera developed low neutralizing titers of 26 and 62, respectively; these mice developed high BA.5 RNA levels in the respiratory tract after challenge (**Figs. 1K-L**). In contrast, the BA.2.12.1- and BA.5-infected sera showed higher neutralization titers of 471 and 375, respectively; these mice developed lower BA.5 RNA levels in the respiratory tracts.

### Antigenic analysis of neutralization titers against variant spikes

Using the neutralizing titer results in **Figs. 1E-H**, we created antigenic maps to visualize the relationships between sera collected from infected mice against SARS-CoV-2 variants and Omicron sublineages (**Fig. 3)**. FFRNT titers were used to position the serum relative to each virus using antigenic cartography (Smith et al., 2004). Each antigenic gridline of the map corresponds to a 2-fold difference in the neutralization titer of a given virus. The cartography illustrated that mouse sera raised to Omicron sublineage variants are antigenically related, with the BA.2 reference being central to the group. Additionally, they formed a new antigenic cluster distinct from the index virus or delta variant (>16-fold titer difference). BA.1-raised mouse antisera exhibited a diverse range in both homologous and heterologous FFRNT_50_ responses, with some sera showing BA.1 specific reactivity and others demonstrating further cross-reactivity to BA.2 and BA.2.12.1 viruses and limited cross-reactivity to BA.4.6 and BA.5 viruses. In contrast, BA.2- and BA.2.12.1-raised mouse antisera demonstrated broader neutralizing responses (≤8-fold reduction) to all Omicron sublineages tested. While BA.5-raised mouse antisera exhibited good cross reactivity to BA.2, BA.2.12.1 and BA.4.6 viruses, they had reduced reactivity to BA.1 virus.

**Figure 3.**
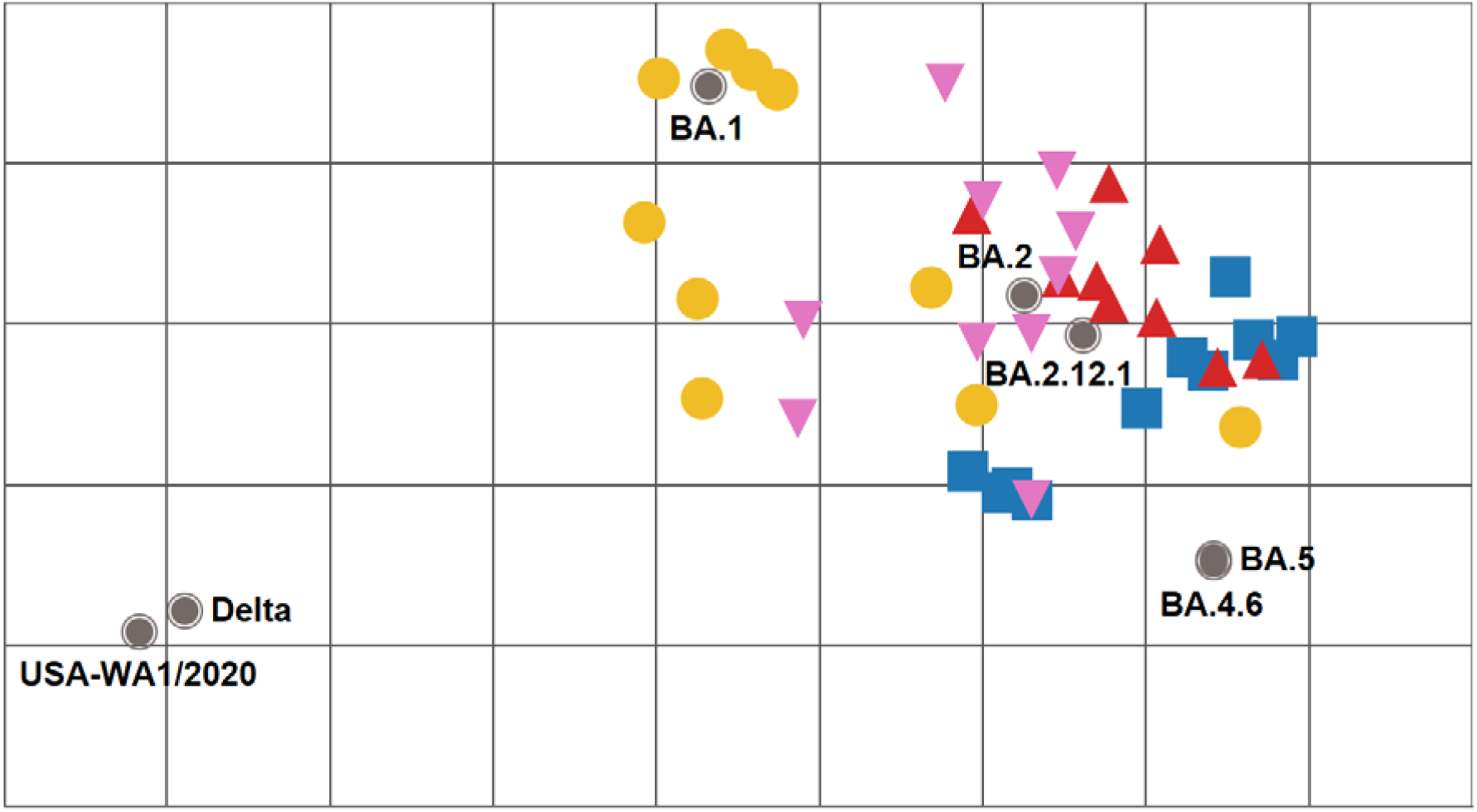
Antigenic cartography of SARS-CoV-2 FFRNT50 data using mouse anti-sera. The antigenic map was prepared according to serum FFTNT_50_ values against mNG reporter SARS-CoV-2 spike variants. The FFTNT_50_ values are presented in **Figures 1E-H**. SARS-CoV-2 variants (gray, filled circle icons) are labeled by name or pango lineage. Individual mouse antisera are color-coded based on the viral inoculum: BA.1 (gold circles), BA.2 (pink triangles), BA.2.12.1 (red triangles), and BA.5 (blue squares). Grid increments indicate a two-fold reduction in FFRNT_50_ titer between two icons. Thus, two grids correspond to 4-fold dilution, three to 8-fold dilution, and so on.

## Discussions

For any emerging SARS-CoV-2 variant, it is important to monitor their viral transmissibility, disease severity, and immune evasion. A systematic analysis of the antigenic relationship among variants provides the scientific basis for future vaccine design. We previously showed distinct cross-neutralization and protection among Alpha, Beta, Gamma, Delta, Epsilon, and Omicron BA.1 variants (Liu et al., 2022c). The current study extends the cross-neutralization analysis to the recently emerged Omicron sublineages, including BA.1, BA.2, BA.2.12.1, BA.4.6, and BA.5. Our antigenic cartography shows that all Omicron sublineages to date are antigenically related to each other and clearly form a new cluster/group from the variants that caused previous global epidemics (*i.e*., index, alpha, delta). The antigenic distance between the index virus and the currently prevalent Omicron BA.5 supports the recommendation for bivalent vaccines containing Omicron variants and suggests that BA.5 should increase the breadth of immunity particularly against decendent lineages or BA.2. Indeed, preliminary clinical results showed that, compared with the original mRNA vaccine, the BA.5-modified bivalent booster elicited more robust neutralization against BA.5 virus (https://www.pfizer.com/news/press-release/press-release-detail/pfizer-and-biontech-announce-positive-early-data-clinical).

For viral replication, the K18-hACE2 mouse data and HAE results consistently suggest the viral fitness in the order of BA.5 ≥ BA.2 ≥ BA.2.12.1 > BA.1. However, no significant difference in weight loss was observed among different sublineage-infected mice. These results indicate that, compared with the initial BA.1, BA.5 has increased viral fitness without increasing disease severity. Our results are in agreement with the clinical observations that (i) BA.1- and BA.5-infected patients develop similar disease severity and (ii) Omicron causes less severe disease than the previous Delta variant (Wolter et al., 2022). The reduced disease severity of Omicron infection may be caused by the lower cell-to-cell fusion activity mediated by the Omicron S protein (Du et al., 2022; Meng et al., 2022). Thus, the increased viral fitness and immune evasion of BA.5 may account for its dominance over other Omicron sublineages in circulation.

Our study has two limitations. First, we have not defined the mutations that determine the differences in viral fitness and/or immune evasion among different Omicron sublineages. These determinants are most likely within the S gene; however, mutations outside the S gene may also contribute to the observed differences. Second, we have not analyzed cell-mediated immunity or non-neutralizing antibodies. These two immune components, together with neutralizing antibodies, protect patients from severe disease and death (Barouch, 2022). Fortunately, after natural infection or vaccination, T cell epitopes are highly preserved in Omicron and other variant S proteins (Redd et al., 2022). Despite the limitations of this study, laboratory investigations, together with real-world vaccine effectiveness results, will continue to guide vaccine strategy and mitigate the public health impact of future variants.

## METHOD

### Ethics statement

Mouse studies were performed following the guidelines for the Care and Use of Laboratory Animals of the University of Texas Medical Branch (UTMB). The protocol (IACUC#: 2103023) was approved by the Institutional Animal Care and Use Committee (IACUC) at UTMB. All the animal operations were performed under anesthesia by isoflurane to minimize animal suffering. All SARS-CoV-2 infections were performed at the BSL-3 facility at UTMB by personnel equipped with powered air-purifying respirators.

### Cells and animals

African green monkey kidney epithelial Vero-E6 cells (laboratory-passaged derivatives from ATCC CRL-1586) were grown in Dulbecco’s modified Eagle’s medium (DMEM; Gibco/Thermo Fisher, Waltham, MA, USA) with 10% fetal bovine serum (FBS; HyClone Laboratories, South Logan, UT) and 1% antibiotic/streptomycin (P/S, Gibco). Vero-E6-TMPRSS2 cells were purchased from SEKISUI XenoTech, LLC (Kansas City, KS) and maintained in 10% fetal bovine serum (FBS; HyClone Laboratories, South Logan, UT) and 1% P/S and 1 mg/ml G418 (Gibco). The primary human airway epithelium (HAE) and medium for culturing HAE were purchased from MatTek Life Science (Ashland, MA, USA). All cells were maintained at 37°C with 5% CO_2_. All cell lines were verified and tested negative for mycoplasma. K18-hACE2 c57BL/6J mice (strain: 2B6.Cg-Tg(K18-ACE2)2Prlmn/J) mice were purchased from the Jackson Laboratory (Bar Harbor, ME). Upon arrival at the ABSL-3 facilities at UTMB, animals were randomized and housed in groups of < 5 per cage in rooms maintained between 68-74°F with 30%–60% humidity and day/night cycles of 12 h intervals (on 6 AM-6 PM). Mice were fed standard chow diets. Female mice aged 8-10 weeks were used for this study.

### Construction of recombinant SARS-CoV-2

The infectious cDNA clones of Omicron sublineages were constructed through mutagenesis of a previously reported cDNA clone of USA-WA1/2020 SARS-CoV-2 (Xie et al., 2021; Xie et al., 2020; Liu et al., 2022a). The Omicron sublineages BA.1 (GISAID EPI_ISL_6640916), BA.2 (GISAID EPI_ISL_11253924.1), BA.2.12.1 (GISAID EPI_ISL_12115772), and BA.5 (GISAID EPI_ISL_11542604) were used as the reference sequences for constructing the infectious cDNA clones. Omicron sublineage BA.1-, BA.2-, BA.2.12.1-, BA.5-spike and Delta-spike mNG SARS-CoV-2s constructed by engineering the complete spike gene from the indicated variants into an infectious cDNA clone of mNG USA-WA1/2020 were reported previously (Kurhade et al., 2022b; Xie et al., 2020). BA.4.6-spike mNG SARS-CoV-2 (GISAID EPI_ISL_15380489) was constructed using the same approach. Supplementary Figure 1 illustrates the spike mutations of Omicron sublineages in this study.

The full-length cDNA was assembled via *in vitro* ligation and used as a template for *in vitro* transcription. The full-length viral RNA was then electroporated into Vero E6-TMPRSS2 cells. On day 2-4 post electroporation, the original P0 virus was harvested from the electroporated cells and propagated for another round on Vero E6 cells to produce the P1 virus. The infectious titer of the P1 virus was quantified by plaque assay on VeroE6-TMPRSS2 cells (recombinant SARS-CoV-2) or by fluorescent focus assay on Vero E6 cells (recombinant SARS-CoV-2 mNG). The P1 viruses were sequenced to ensure no undesired mutations. The P1 virus was used for all the experiments performed in this study.

### RNA extraction and Sanger sequencing

Culture supernatants were mixed with a five-fold volume of TRIzol™ LS Reagent (ThermoFisher Scientific). Viral RNAs were extracted by using the Direct-zol-96 MagBead RNA kit (ZYMO RESEARCH) in the KingFisher Apex instrument (ThermoFisher Scientific) according to the manufacturer’s instructions. The extracted RNAs were eluted in 50 μl nuclease-free water. The cDNA fragments spanning the genome were amplified using SuperScript™ IV One-Step RT-PCR System (ThermoFisher Scientific, Waltham, MA) according to a previously reported protocol (Xie et al., 2021; Xie et al., 2020). The resulting fragments were gel purified and sent for Sanger sequencing at GENEWIZ.

### Mouse study

8-10-week-old female K18-hACE2 mice were infected intranasally (i.n.) with infectious cDNA clone derived SARS-CoV-2 BA.1, BA.2, BA.2.12.1 and BA.5 (10^4^ PFU, diluted in 25 μl DPBS). DBPS-inoculated animals were used as controls. On day 2 post-infection, lung (cranial lobe) and trachea samples were collected from a subset of euthanized animals. The animals were monitored for signs of disease and weighed daily for 21 days after the primary infection. On day 21 post-infection, animals were anesthetized, and blood samples were collected retro-orbitally. Sera were isolated and stored at −80°C before use. On day 38 post-infection, all animals were re-challenged with infectious cDNA clone-derived BA.5 (10^4^ PFU, diluted in 25 μl DPBS). Two days after the challenge, all animals were euthanized. Mouse lung cranial lobes and tracheas were harvested in 2-ml tubes that were prefilled with 1 ml DPBS; the other lobes of the right lung and nasal turbinates were kept in 2-ml tubes that were prefilled with 500 μl Trizol reagent (ThermoFisher Scientific). All samples were stored at −80°C before analysis. Tissue samples were homogenized for 60 seconds using MagNA Lyser (Roche) with settings of 6000 rpm and clarified by centrifugation at 12,000 g for 5 min before being used for plaque assay and RNA extraction.

### Plaque assay

Approximately 1.0 × 10^6^ Vero E6-TMPRSS2 cells were seeded to each well of 6-well plates and cultured at 37 °C, 5% CO_2_ for 16 h. Virus samples were serially diluted in DMEM with 2% FBS and 200 μl diluted viruses were transferred onto the cell monolayers. The viruses were incubated in the cells at 37 °C with 5%CO_2_ for 1 h. After the incubation, the inoculum was removed. Two ml of overlay medium containing DMEM with 2%FBS, 1% penicillin/streptomycin, and 1% Seaplaque agarose (Lonza, Walkersville, MD) were added to each well of the infected cells. After incubation at 37°C with 5% CO_2_ for 2 days, the cells were stained by adding 2 ml of overlay medium supplemented with neutral red (SigmaAldrich, St. Louis, MO). Plaques were counted on a lightbox.

### Quantitative real-time RT-PCR

RNA copies of SARS-CoV-2 samples and the mouse housekeeping gene GAPDH were determined by quantitative real-time RT-PCR (RT-qPCR) by using the iTaq SYBR Green One-Step Kit (Bio-Rad) on the QuantStudio™ 7 Flex system (ThermoFisher Scientific) with the following settings: (1) 50°C, 10 min; 95°C, 5 min; (2) 95°C, 15 s; 60°C, 30 s; 40 cycles; (3) 95°C, 15 s; 60°C to 95°C, increment 0.5°C, 0.05 s. The primer pairs for the SARS-CoV-2 N gene and mouse GAPDH gene are the following: 2019-nCoV_N2-F (5’-TTACAAACATTGGCCGCAAA-3’) and 2019-nCoV_N2-R (5’-GCGCGACATTCCGAAGAA-3’); M_GAPDH-F (5’-AGGTCGGTGTGAACGGATTTG-3’) and M_GAPDH-R(5’-TGTAGACCATGTAGTTGAGGTCA-3’). The relative viral RNA levels (N-gene to GAPDH ratio) were obtained by normalizing the CT values of the N gene to those of the GAPDH in each sample. The cutoff of the N-gene to GAPDH ratio was calculated as the mean plus two times of standard deviation from uninfected mouse samples. The log10 values of the cutoffs of N-gene/GAPDH in the lungs and turbinates are −4.92 and −4.18, respectively.

### Fluorescent focus reduction neutralization test

A fluorescent focus reduction neutralization test (FFRNT) was performed to measure the neutralization titers of sera against USA-WA1/2020, Delta-spike, *BA.1-, BA.2-, BA.2.12.1-, BA.4.6-, and BA.5-spike* mNG SARS-CoV-2. The FFRNT protocol was reported previously (Zou et al., 2022b). Vero E6 cells were seeded onto 96-well plates with 2.5×10^4^ cells per well (Greiner Bio-one™) and incubated overnight. On the next day, mouse sera were heat-inactivated at 56°C for 30 min before assay. Each serum was 2-fold serially diluted in a culture medium and mixed with 100-150 focus-forming units of mNG SARS-CoV-2. The final serum dilution ranged from 1:20 to 1:20,480. After incubation at 37°C for 1 h, the serum-virus mixtures were loaded onto the pre-seeded Vero E6 cell monolayer in 96-well plates. After 1 h infection, the inoculum was removed and 100 μl of overlay medium containing 0.8% methylcellulose was added to each well. After incubating the plates at 37°C for 16 h, raw images of mNG foci were acquired using Cytation™ 7 (BioTek) armed with 2.5× FL Zeiss objective with a wide field of view and processed using the software settings (GFP [469,525] threshold 4000, object selection size 50-1000 μm). The fluorescent mNG foci were counted in each well and normalized to the non-serum-treated controls to calculate the relative infectivities. The FFRNT_50_ value was defined as the minimal serum dilution to suppress >50% of fluorescent foci. The neutralization titer of each serum was determined in duplicate assays, and the geometric mean was taken.

### Antigenic Cartography

FFRNT_50_ titer responses were transformed into a two-dimensional (2D) antigenic map over 1000 iterations with the ACMACS-API software suite (version acmacs-c2-20161026-0717 and i19 build host) (Smith et al., 2004). Output data matrices with FFRNT_50_ trends mapped to 2D X/Y coordinates were rendered into antigenic landscapes and annotated in Tableau Desktop (version 2021.1.1). All applied code-sets are available from the authors upon request.

### Competition experiment

For the competition on HAE, two viruses were mixed with equal PFU based on the titers as determined on VeroE6-TMPRSS2 cells. The inoculum was prepared in DPBS with 20% culture medium containing virus mixture at a final concentration of 1×10^6^ PFU/ml. An aliquot of the inoculum was also stored for determining the ratio of input viruses. Before infection, the HAE culture was incubated in 300 μl of DPBS per well at 37°C. After removing the DPBS, 200 μl of inoculum per well was added to each HAE well at the apical side. After 2 h-incubation at 37°C 5% CO_2_, the inoculum was removed, and cells were washed with DPBS three times to remove unbound viruses. At each sample collection time point, 300 μl of DPBS was added to each well at the apical side, and cells were incubated at 37°C 5% CO_2_ for 30 min. The DPBS washes containing viruses were then harvested into 2-ml tubes. All samples were stored in a −80°C freezer before use.

Upon analysis, samples were thawed at room temperature. 100 μl of each sample was used for Trizol LS extraction using the protocol described above. The cDNA fragment (corresponding to codon positions 411-479 of BA.1 spike) containing the unique mutations of each variant was amplified using SuperScript™ IV One-Step RT-PCR System with primer pairs: BA.1-2-4-F(5’-CAAACTGGAAATATTGCTGA-3’) and BA.1-2-4R (5’-CCATTACAAGGTTTGTTACC-3’). The amplicons were then gel-purified and sent for Illumina next-generation sequencing (NGS) at the sequencing core facility at UTMB.

### Statistical Analysis

Numeric data are presented as means ± standard deviations or geometric mean ± 95% confident intervals as indicated in each figure panel. Data in Figure 1 C-D and I-L were initially log(base-10) transformed to increase the normal distribution and one-way ANOVA with Tukey’s multiple comparisons test was applied to assess the statistical significance between each group shown in. The Wilcoxon matched-pairs signed-rank test was performed for group comparison of GMTs of mouse serum in Figure 1 E-H. Simple linear regression was performed to analyze the statistical significance of the RNA ratio at a given time-point versus the input RNA ratio for the competition experiment in Figure 2. All statistical analysis was performed using the software Prism 9 (GraphPad).

## Acknowledgments

We thank colleagues at the University of Texas Medical Branch (UTMB) for helpful discussions and Samuel S. Shepard and Reina Chau for their contributions in *operationalizing ACMACS for high-throughput and portable usage*. This research was partly funded by the Centers for Disease Control and Prevention. P.-Y.S. was supported by NIH contract HHSN272201600013C, and awards from the Sealy & Smith Foundation, the Kleberg Foundation, the John S. Dunn Foundation, the Amon G. Carter Foundation, the Summerfield Robert Foundation, and Edith and Robert Zinn. SSS is affiliated with the Centers for Disease Control and Prevention (CDC). RC is a CDC contractor with General Dynamics Information Technologies, Inc. Use of trade names is for identification only and does not imply endorsement by the US Centers for Disease Control and Prevention or the US Department of Health and Human Services. The findings and conclusions in this report are those of the authors and do not necessarily represent the official position of the US Centers for Disease Control and Prevention.

## Author contributions

Conceptualization, X.X., P.-Y.S.; Methodology, H.X., J.Y., B.K., C.B., J.C., C.K., J.Z., S.G.W., B.R.M., R.K., C.T.D., B.Z., D.E.W., X.X., P.-Y.S; Investigation, H.X., J.Y., B.K., C.B., J.C., C.K., J.Z., S.G.W., B.R.M., R.K., C.T.D., B.Z., D.E.W., X.X., P.-Y.S; Resources, X.X., P.-Y.S; Data Curation, CDC colleagues, X.X.; Writing-Original Draft, B.R.M., R.K., D.E.W., X.X., P.-Y.S; Writing-Review & Editing, R.K., D.E.W., X.X., P.-Y.S.; Supervision, H.X., B.K.K, R.K., D.E.W., X.X., P.-Y.S.; Funding Acquisition, P.-Y.S.

---

**Supplementary Figure 1.**
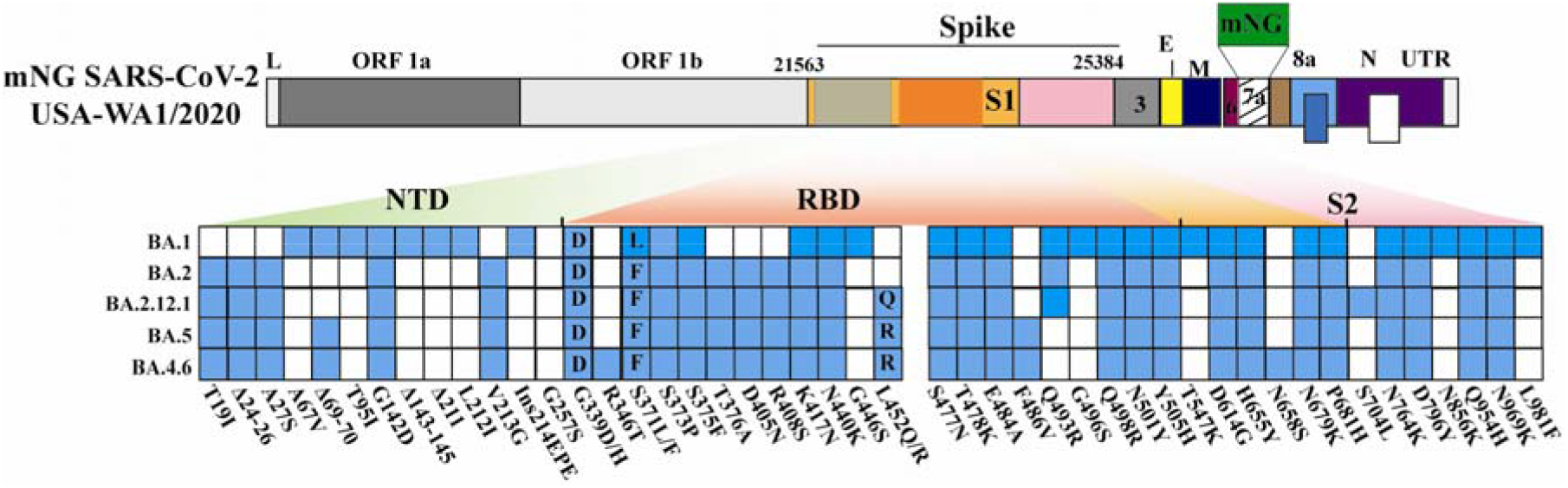
Diagram of spike mutations of Omicron sublineages. mNG USA-WA1/2020 SARS-CoV-2 was used to engineer Omicron-spike SARS-CoV-2s. The mNG reporter gene was engineered at the open-reading-frame-7 (ORF7) of the USA-WA1/2020 genome.1 Amino acid mutations, deletions (Δ), and insertions (Ins) are indicated for variant spikes in reference to the USA-WA1/2020 spike. L: leader sequence; ORF: open reading frame; NTD: N-terminal domain of S1; RBD: receptor binding domain of S1; S: spike glycoprotein; S1: N-terminal furin cleavage fragment of S; S2: C-terminal furin cleavage fragment of S; E: envelope protein; M: membrane protein; N: nucleoprotein; UTR: untranslated region.

